# Circadian clock function does not require the histone methyltransferase MLL3

**DOI:** 10.1101/2021.12.10.472092

**Authors:** Matthew Baxter, Toryn Poolman, Peter Cunningham, Louise Hunter, Maria Voronkov, Gareth B. Kitchen, Laurence Goosey, Nicola Begley, Danielle Kay, Abby Hespe, Robert Maidstone, Andrew S.I. Loudon, David W. Ray

**Affiliations:** NIHR Oxford Biomedical Research Centre, John Radcliffe Hospital, Oxford, UK and Oxford Centre for Diabetes, Endocrinology and Metabolism, University of Oxford, Oxford, OX3 9DU, UK; Centre for Biological Timing, Faculty of Biology, Medicine and Health, University of Manchester, Manchester, M13 9PT, UK

**Keywords:** Circadian, clock, epigenetics, histone, methyltransferase, MLL3, KMT2C, inflammation

## Abstract

The circadian clock controls the physiological function of tissues through the regulation of thousands of genes in a cell-type specific manner. The core cellular circadian clock is a transcription-translation negative feedback loop, which can recruit epigenetic regulators to facilitate temporal control of gene expression. Histone methyltransferase, mixed lineage leukemia gene 3 (MLL3) was reported to be required for maintenance of circadian oscillations in cultured cells. Here, we test the role of MLL3 in circadian organisation in whole animals. Using mice expressing catalytically inactive MLL3, we show that MLL3 methyltransferase activity is in fact not required for circadian oscillations *in vitro* in a range of tissues, nor for maintenance of circadian behavioural rhythms *in vivo*. In contrast to a previous report, loss of MLL3-dependent methylation did not affect global levels of H3K4 methylation in liver, indicating substantial compensation from other methyltransferases. Further, we found little evidence of genomic repositioning of H3K4me3 marks. We did, however, observe repositioning of H3K4me1 from intronic regions to intergenic regions and gene promoters, however there were no changes in H3K4me1 mark abundance around core circadian clock genes. Output functions of the circadian clock, such as control of inflammation, were largely intact in MLL3-methyltransferase deficient mice, although some gene specific changes were observed, with sexually dimorphic loss of circadian regulation of specific cytokines. Taken together, these observations call for a major reassessment of the inter-relationship between the circadian clock and MLL3-directed histone methylation, and a deeper examination of other epigenetic mechanisms which may facilitate circadian clock function.

**Significance statement:** A highly cited paper published in PNAS previously reported an essential role for the histone methyltransferase MLL3 in maintaining circadian oscillations in cultured cells.

We tested the role of MLL3 *in vivo* and in primary tissues showing that MLL3 in fact plays no role in organising the core circadian clock, and has no functional impact on whole animal circadian behaviour. However, in further analysis, we newly discover a role for MLL3 in conferring circadian control to components of the inflammatory response, doing so in a sexually dimorphic manner. As the MLL family of histone methyltransferases are being targeted by pharmaceuticals for cancer, it is important to understand how methyltransferases may be driving circadian rhythms in gene expression.

## Introduction

The circadian clock drives the expression of thousands of genes in diverse tissues, conferring on them oscillating activity. The molecular clockwork consists of multiple transcriptional regulators which operate in a transcription-translation feedback loop (TTFL), with a period of approximately 24 hours. Core clock components include BMAL1 and CLOCK, which form a heterodimer, capable of transactivating numerous target genes by binding to conserved E-box elements. Amongst these targets are the PER and CRY genes, which form a transcriptional repressive complex to inhibit BMAL1/CLOCK transactivation. This completes the feedback loop. In addition, a second negative feedback loop arises from BMAL1/CLOCK transactivation of the NR1D1 orphan nuclear receptor. This binds to RORE elements on genomic DNA and recruits NCOR/HDAC3 repressor complexes to inhibit expression of BMAL1. NR1D1 binds the same elements as RORa, a transactivator. In contrast to the high amplitude NR1D1 circadian oscillation RORa expression is stable through time.

In addition to the well-defined role of transcription factors in driving circadian oscillations in gene expression there is evidence for dynamic changes in histone modifications associated with many circadian-target genes, including Dbp, Per1, and Per2 (Etchegaray et al., 2003; Kwapis et al., 2018; Naruse et al., 2004; Stratmann et al., 2012). Regulation of histone modifications are likely to work in concert with the core clock TTFL, in order to confer specificity of transcriptomic rhythmicity to different tissues and cell types. To date, most research has focussed on histone methylation and acetylation, and various marks, especially H3K4me1, H3K4me3, and H3K27ac are found to show marked changes in abundance through circadian time series (Koike et al., 2012). Histone methyltransferases of the mixed lineage leukemia (MLL) family have emerged as likely effectors and, indeed, the expression of MLL1 and MLL3 may be clock controlled in specific tissues (Katada and Sassone-Corsi, 2010; Valekunja et al., 2013). MLL3 was originally thought to induce H3K4me3 at the promoter regions of genes, however more detailed investigations suggest it mainly contributes to H3K4me1 at enhancer elements (Herz et al., 2012; Jozwik et al., 2016; Lee et al., 2008).

MLL3 has been proposed as an epigenetic regulator required for the molecular circadian clock to function (Valekunja et al., 2013). Valekunja et al reported MLL3 to be a clock-controlled factor that directly or indirectly modulates over 100 epigenetically targeted circadian output genes in the mouse liver. They also showed almost complete loss of genome-wide H3K4me3 deposition in MLL3 methyltransferase-deficient mouse embryonic fibroblasts (MEFs). Furthermore, Valekunja et al claimed that catalytic inactivation of MLL3 severely compromises the oscillation of core clock genes, including *Bmal1, Rev-erbα*, and *Per2*, and concluded that lack of MLL3 methyltransferase activity controls both “core” and “output” clock genes (Valekunja et al., 2013).

Here, we sought to extend the findings of Valekunja et al to investigate the physiological role of MLL3-dependent methylation in maintaining circadian organisation, and temporal programming of tissue responses. Surprisingly, and in contrast to the previous report, we found that MLL3 methyltransferase activity was not required for core circadian function. Furthermore, we found almost no difference between genome-wide H3K4me3 levels in MLL3 methyltransferase deficient mouse livers compared to wild-type littermate controls. Similarly, there was only subtle repositioning of genome wide H3K4me1 deposition in liver between the two genotypes. Lastly, we examined the role of MLL3 in models of circadian output, namely, gating of immune function. Here, we found that MLL3 was required for subtle, and gene-specific time of day variation of innate immunity. Taken together, we refute a central role for MLL3 in the operation of the molecular circadian clock, but discover a subtle, but distinct, role for MLL3 in coupling inflammatory signalling to the circadian clock.

## Results

### MLL3 Delta mice show normal behavioural circadian rhythms

MLL3 functions as part of a multi-protein complex known as ASCOM, which also contains RbBP5, ASH2L, WDR5, UTX, and other additional proteins (S. Lee, Roeder, & Lee, 2009). ASCOM may contain either MLL3 or MLL4 as the active H3K4 methyltransferase, whilst UTX demethylates H3K27. ASCOM may also recruit CBP/p300 to acetylate H3K27, further facilitating gene transcription (Lee et al., 2006, 2001). To explore the biology of MLL3-mediated methylation specifically, we used a strain of mice (MLL3 Delta) which lacked the catalytic domain of the MLL3 enzyme due to the in-frame deletion of exons 57-58 (Lee et al., 2006). This mutation renders MLL3 unable to catalyse methylation of H3K4, whilst preserving structural functions. The previous reports of a striking circadian role for MLL3 were defined using isolated MEF cells from these same MLL3 Delta mice in culture (Valekunja et al., 2013). Here, we bred MLL3 Delta mice on a mixed C57Bl6/129sv genetic background, since this mutation is embryo-lethal on a pure C57Bl6 background (Fig S1A) (Lee et al., 2008). Using this breeding strategy, we were able to generate viable mice, homozygous for the MLL3 catalytic domain mutation (henceforth referred to as MLL3 Delta mice), and wild-type littermate controls. We observed fewer live-born MLL3 Delta animals, in line with previous observations (Lee et al., 2008), suggesting that MLL3 catalytic deficiency imposes a toll in animal development (Fig S1B). We considered that there may be compensation of other methyltransferases seen in survivors which may impact our model, but saw no such changes when we measured expression of the related methyltransferases MLL1, 2, and 4 (Fig S1C). We were also able to confirm that MLL3 expression levels were unaffected, despite the expected in-frame deletion of exons 57 and 58 (Fig S1C).

The mice were left to free-run in constant darkness (Fig 1A). Free-running period was slightly but non-significantly shorter in MLL3 Delta mice compared to littermate controls and less than 24 hours for both gneotypes (Fig 1B), consistent with previously published observations of mice on a C57Bl6 genetic background. In further analyses we measured activity during the light and dark phases (Fig 1C) and found no genotype differences. We next tested phase shifting responses, and exposed mice to a 6-hour phase advance (Fig 1D). There was no significant difference between the MLL3 Delta mice and the WT littermate controls in the time taken to re-set (Fig 1E). These findings were strikingly different from predictions based on the *in vitro* analysis of MLL3 circadian function (Valekunja et al., 2013).

**Figure 1.**
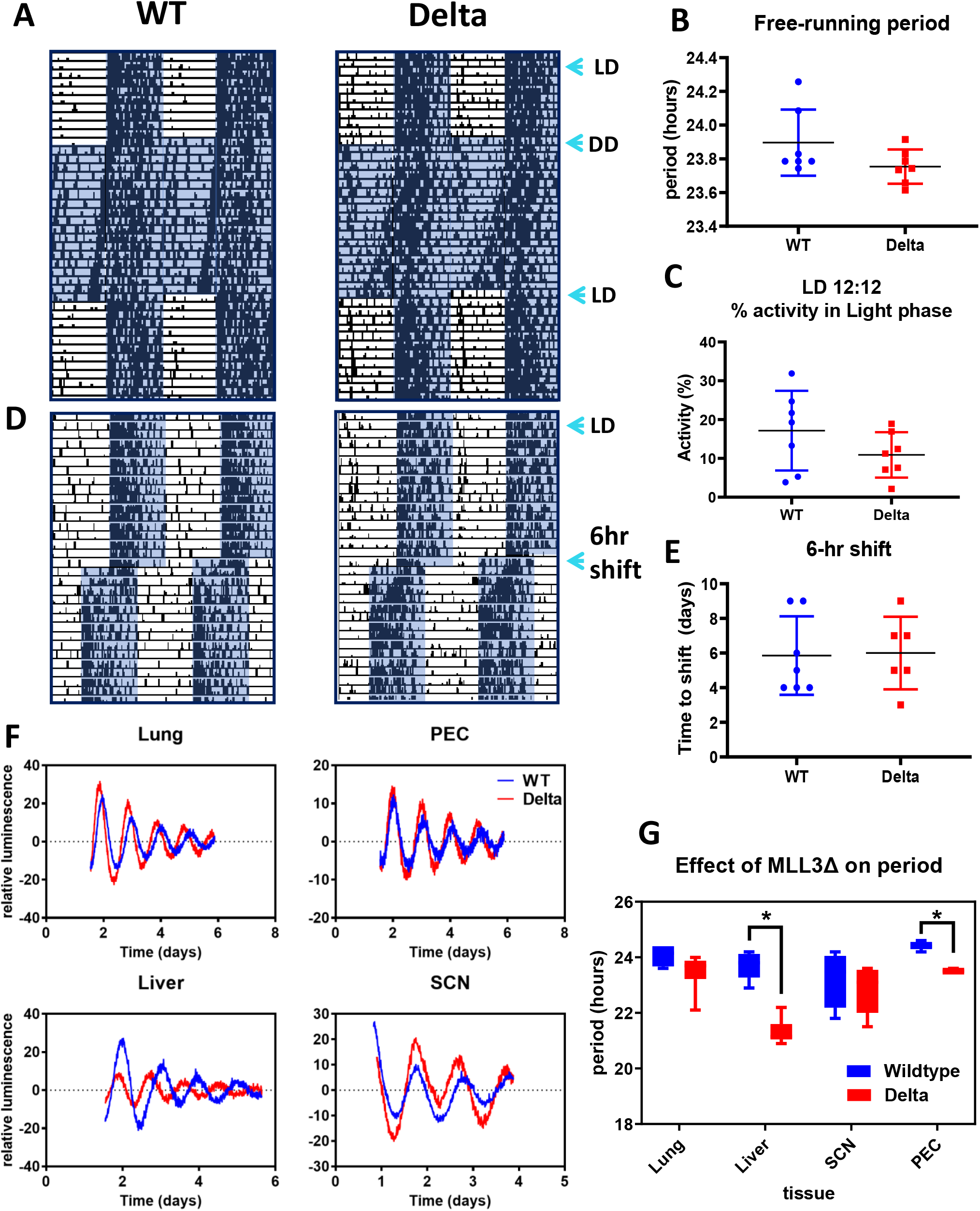
Circadian rhythms persist in the absence of MLL3 methyltransferase function. **A**. Mice expressing global MLL3 Delta and littermate wild-type controls were placed in light control cabinets in 12:12 light dark (L:D) cycles for 2 weeks to acclimatize. Mice were then exposed to constant dark for 2 weeks and allowed to “free-run”, before returning to a 12:12 L:D exposure. Mouse activity was monitored using beam-break technology. **B**. The free-running period was calculated during two weeks of constant darkness. **C**. The percentage of activity in the light phase of each animal was quantified during one week in 12:12 L:D conditions. **D**. After another 2 week acclimatization period to 12:12 L:D conditions, mice were exposed to a 6-hr phase advance light shift. **E**. The time-to-shift and re-entrain with the phase advance light pattern was quantified. **F**. Lung, liver, suprachiasmatic nuclei (SCN), and peritoneal macrophage (PEC) tissue explants from MLL3 Delta mice and wild-type littermate controls on a Per2::luc genetic background were placed in a lumicycle to observe circadian oscillations in PER2 expression. **G**. The circadian period in each of the tissues was quantified and compared between the two genotypes (n=3-5).

As previous reports had detected a major role for MLL3 in isolated cells in culture we considered that the minimal effects observed *in vivo* may result from masking phenomena arising from a dominant central pacemaker. Therefore, we crossed the MLL3 delta mice onto the Per2::Luc genetic background. We isolated four tissue types, namely liver, lung, suprachiasmatic nucleus (SCN) and peritoneal macrophages (PEC) and monitored these cells, and tissue explants in culture through time, monitoring light production in a lumicycle (Fig 1F,G). In marked constrast to the earlier report (Valekunja et al., 2013), all isolated tissues showed strong, sustained *in vitro* circadian oscillations, allowing detailed analysis of waveforms. SCN and lung explants showed no significant genotype difference. In contrast, both liver and PECs had a significantly shorter circadian period in MLL3-delta derived tissues (Fig 1G).

### ChIP-seq reveals no global change in H3K4me3 levels in liver

To investigate further possible mechanisms underpinning the altered circadian phenotype seen in hepatic tissue, we performed ChIP-Seq for the H3K4me3 mark in liver from MLL3 Delta animals and wild-type littermate controls. We selected Zeitgeber-time 18 (ZT18), as this timepoint had previously been proposed as the peak for H3K4me3 levels and genomic MLL3 association (Koike et al., 2012; Valekunja et al., 2013). Strikingly, we observed remarkably few differences between MLL3 Delta animals and wild-type littermate controls. In contrast to previous findings (Valekunja et al., 2013), we identified no impact of MLL3 Delta on global H3K4me3 levels (Fig 2A). We further noted there was no observable difference in H3K4me3 levels at the promoter of any core clock gene, in agreement with the intact circadian oscillations observed in mouse behaviour and isolated tissues (Fig 2B). In total, 36,079 H3K4me3 peaks were identified in wildtype mice, of which 31,234 (87%) were also identified in the MLL3 Delta mice, highlighting the remarkable similarity between the genotypes (Fig 2C). Indeed, princicpal component analysis did not separate the animals by genotype, only by sex (Fig 2 Supp B)

**Figure 2.**
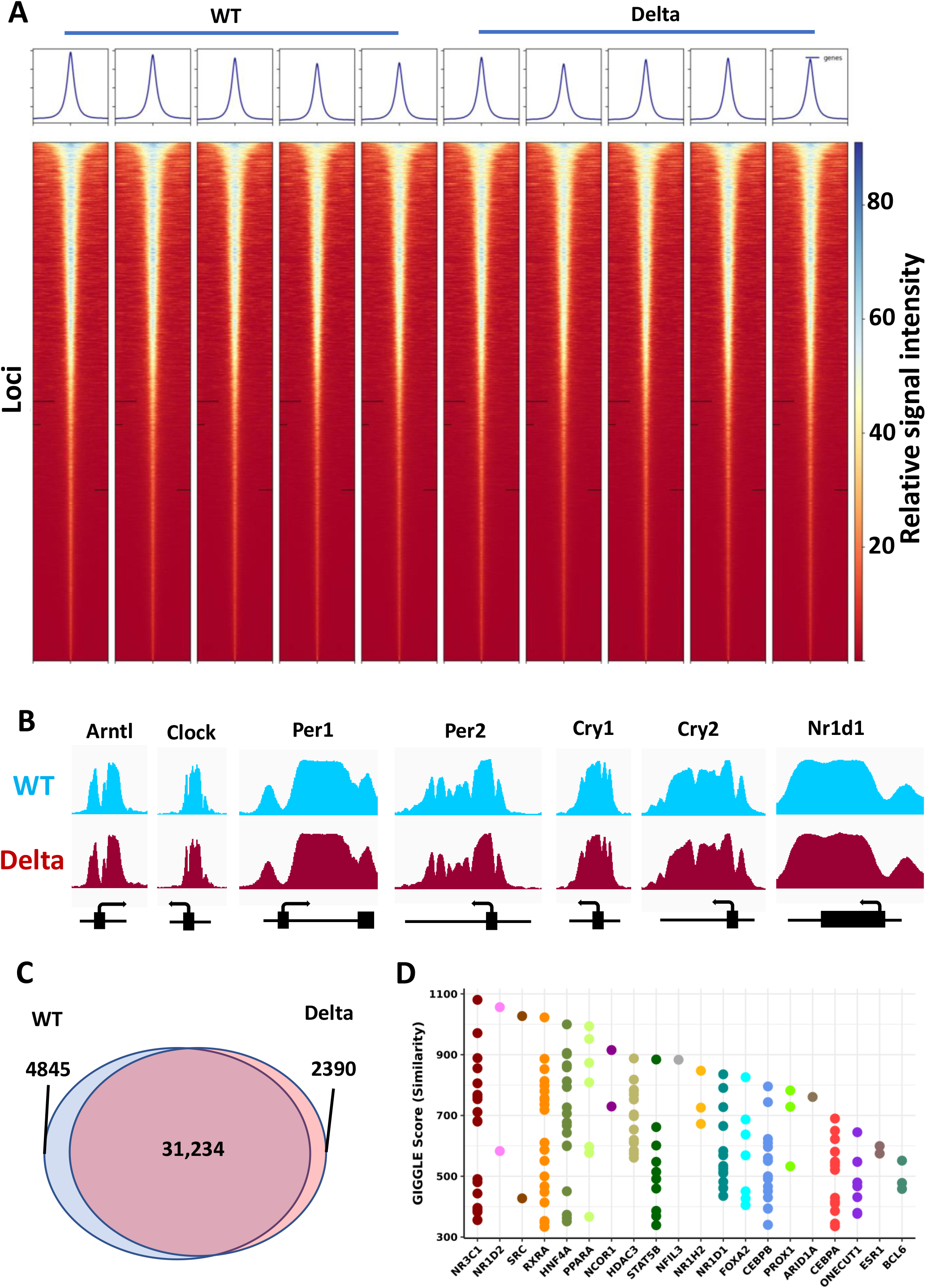
H3K4me3 ChIP-seq analysis of liver tissue from MLL3 Delta mice and WT littermate controls. **A**. Individual heatmaps of MACS2-called H3K4me3 peak intensities from n=5 Delta and n=5 control WT mice were plotted. 5kb either side of the peak summit is shown. **B**. MACS2 derived Bedgraph files for all samples within a group were converted to TDF files to view ChIP-seq gene tracks in IGV. Gene tracks across the promoter and TSS of core clock genes are shown for the WT and MLL3 Delta genotypes.**C**. MACS2 was also used to call peaks from the BAM files. The Venn diagram shows the total number of overlapping and non-overlapping peaks called within the WT and Delta samples. **D**. Peaks which were found to be unique to the WT condition were submitted for Giggle score analysis to assess similarity with other published ChIP-seq factor binding datasets.

### H3K4me3 does not explain changes in period

As we observed a significant shortening of period in the MLL3 Delta liver explants we looked for a potential mechanism involving MLL3 catalytic activity in the H3K4me3 ChIP-seq data. To investigate this we identified the H3K4me3 peaks which were unique to the wild-types or MLL3 Deltas, and also located at a canonical gene promoter, and compared these genes with those listed in the “Circadian rhythm genes” GO term (Fig 2. Supp A). Only two “circadian genes” which lost promoter H3K4me3 peaks were identified, *Drd1* and *Slc6a4*, and one “circadian gene” gained a H3K4me3 peak in the MLL3 Deltas, *Nrg1*. None of these three genes are known to have a role in determining core clock period, nor are they typically well expressed in liver. Therefore, the altered period length seen in MLL3 Delta liver explants (Figure 1F) is not likely to be due to altered H3K4me3 deposition affecting promoter function of the core clock.

To take an unbiased view of the subtle changes in H3K4me3 deposition between the two genotypes, we performed GIGGLE analysis on the loci of the unique wild-type peaks (Zheng et al., 2019). The wild-type unique peaks shared genomic loci with well-characterised MLL3 recruiting transcription factors including the glucocorticoid receptor (NR3C1), and interestingly, the key circadian output regulator REVERBb (NR1D2) (Fig 2D). In addition, there was enrichment for other circadian, or circadian output factors, including PPARA, STAT5B, NFIL3, REVERBa (NR1D1), and NCOR1 (Caratti et al., 2018; Eckel-Mahan et al., 2013; Pariollaud et al., 2018; Quagliarini et al., 2019; Wang et al., 2017; Yin and Lazar, 2005). These overlaps hint that MLL3 enzymatic activity may play a role in tuning circadian output functions, without contributing to the core circadian oscillator.

### Minor repositioning of H3K4me1 in MLL3 Delta liver

Recent studies have shown that MLL3 exhibits higher activity depositing monomethyl than trimethyl marks at H3K4 (Dorighi et al., 2017; Herz et al., 2012; Hu et al., 2013; Local et al., 2018). Mono-methylation of H3K4 denotes enhancers and deposition of this mark has also been linked to circadian transcription (Koike et al., 2012). To pursue the molecular basis of the altered circadian period length seen in isolated liver tissue we then moved to profile changes in H3K4me1 in livers from the two genotypes (Fig 3). Similarly to the H3K4me3 dataset, there was no gross change in the global levels of H3K4me1 marks across the genome (Fig 3A). Similarly, no differences were found in H3K4me1 deposition across circadian genes (Fig 3B). However, some subtle differences were identified between the two genotypes. In total, 80,116 H3K4me1 peaks were identified in the wildtype, of which 68,844 were also found in the MLL3 Delta mice (Fig 3C). Furthermore, 9,744 peaks were found only in the MLL3 Deltas. Interestingly, there appeared to be a subtle, but distinct, redistribution of peaks from introns to intergenic and promoter regions in the MLL3 Delta mice (Fig 3D). We examined reads arising from the MLL3 gene locus, and, as a useful additional control for the efficiency of gene recombination, we saw a complete absence of reads from the targeted MLL3 exons, 57 and 58 (Fig S3A).

**Figure 3.**
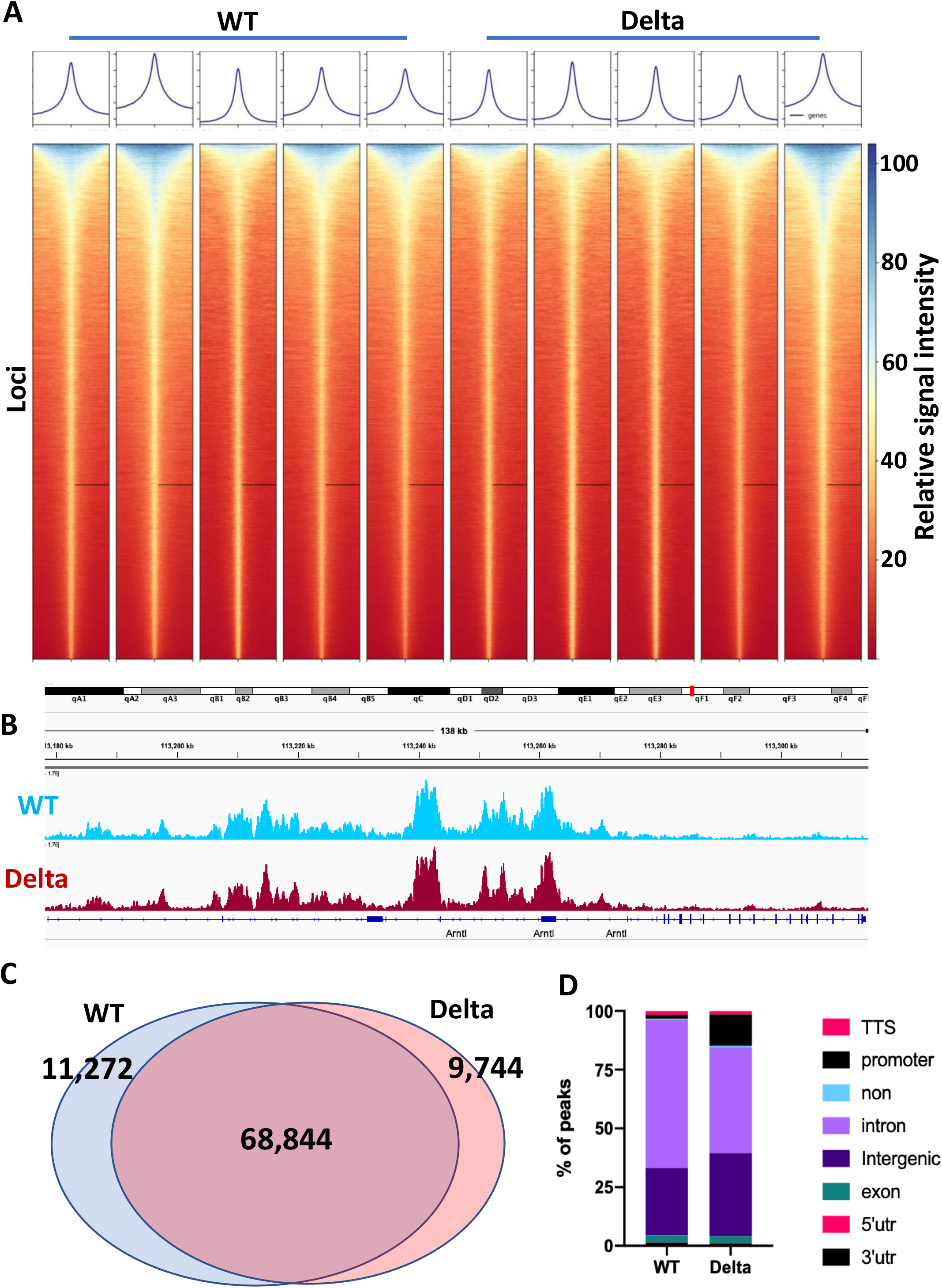
H3K4me1 ChIP-seq analysis of liver tissue from MLL3 Delta mice and WT littermate controls. **A**. Individual heatmaps of MACS2-called H3K4me1 peak intensities from n=5 Delta and n=5 control WT mice were plotted. 5kb either side of the peak summit is shown. **B**. H3K4me1 gene tracks for WT and Delta conditions are shown across the *Bmal1* (*Arntl*) locus. **C**. Venn diagram showing the overlap of MACS-called broad peaks in the WT and Delta conditions. **D**. The relative proportion of peaks across different types of genomic regions was quantified.

Analysis of the genotype unique peaks by GIGGLE revealed a surprising overlap with the variant histone H2AZ at gained peaks in the MLL3 delta, but we did not see enrichment for circadian transcription factors in either condition (Fig 3 Supp B&C).

### Macrophage-specific deletion of MLL3 catalytic region does not affect circadian period

The isolated SCN appeared to show no genotype-dependent circadian phenotype, and its operation *in vivo* may be masking the role of MLL3 on clock function in peripheral cells. Since macrophage cells did show significant period-shortening in MLL3 Delta mice, we explored this further, and targeted MLL3 deletion to specific cells using cell-type specific cre recombinase (Lysozyme Cre) crossed to a floxed allele of MLL3 to target macrophage cells. Surprisingly, and in contrast to macrophages recovered from the global MLL3 Delta animals, the peritoneal macrophages showed no genotype differences (Fig 4A).

**Figure 4.**
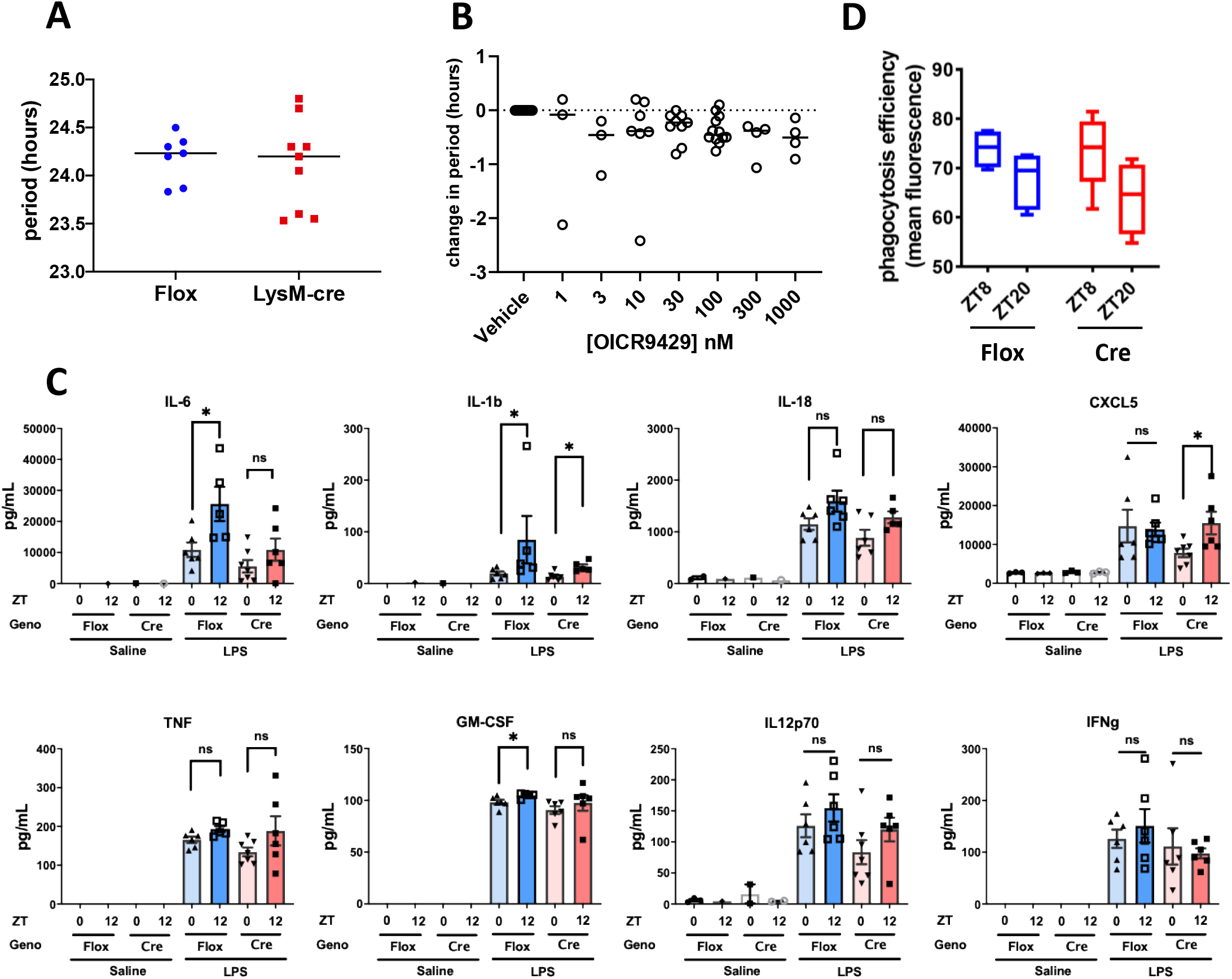
The role of MLL3 in the circadian function of macrophages. **A**. PECs were isolated from MLL3^Delta flox/Delta flox^.LysMcre^het^ mice and MLL3^Delta flox/Delta flox^ littermate controls, and placed into a lumicycle. Circadian oscillations in Per2::luc were measured for 3 days and period was calculated. Each data point represents the average of 3 technical replicates from an individual animal. **B**. PECs were isolated from Per2::luc mice and plated in 96 well plates before treatment with varying concentrations of selective WDR5 inhibitor OICR9429 or vehicle control (0.1% DMSO). Oscillations in Per2::luc were measured using a Clariostar plate reader and period was calculated from 24-72 hours after plating. **C**. MLL3^Delta flox/Delta flox^ LysMcre^het^ mice and MLL3^Delta flox/Delta flox^ littermate controls, were exposed to I.P. LPS at ZT0 or ZT12. 6 hours after exposure cytokine levels in blood plasma were measured by ELISA or Magpix. **D**. PECs were isolated from MLL3^Delta flox/Delta flox^ LysMcre^het^ mice and MLL3^Delta flox/Delta flox^ littermate controls at ZT8 or ZT20 and plated with pHrodo-Staph. Aureus bioparticles. Efficiency of phagocytosis was measured by flow cytometry (n=4-6).

While MLL3 drives the active chromatin H3K4 methyl marks, another enzyme in the COMPASS complex, UTX, removes H3K27me3 marks. We therefore speculated that another histone methyltransferase, EZH2, which deposits H3K27me3, may have a role in circadian function in macrophages. EZH2 has been reported to affect circadian function, and we have shown has impacts on innate immunity, and macrophage function (Kitchen et al., 2021; Zhong et al., 2018). Therefore, we analysed isolated peritoneal macrophages from the *CX3CR1-cre*.*EZH2*^*f/f*^ animals, compared to the cre-negative littermates (Fig S4A). Here again, we did not detect any circadian phenotype, despite the earlier report of a strong circadian phenotype in zebrafish (Zhong et al., 2018). We also analysed liver tissue from a liver-specific EZH2 knockout mouse, on the basis that isolated liver tissue from the MLL3 disrupted animals showed a circadian phenotype, using a tamoxifen-directed deletion strategy to reduce confounding by genetic compensation. However, again, no circadian phenotype was seen (Fig S4B). Therefore, EZH2 did not contribute to circadian organisation in our models.

A degree of functional redundancy between MLL3 and MLL4 within the COMPASS complex has been previously reported (Lee et al., 2008, 2009, 2006). It is possible that this functional redundancy may mask an important role for COMPASS-directed H3K4 methylation in circadian cycling. Therefore we tested an inhibitor of WDR5, which is the common structural platform of the COMPASS complex facilitating the interaction of the methyltransferase enzyme with the histone substrate (Dou et al., 2006). However, there was no observed dose-dependent change in circadian period in isolated macrophages treated with WDR5 antagonist, OICR9429 (Fig 4B).

Taken together these observations may indicate that the differences in period observed in the global MLL3 Delta mice, may not be directly due to changes in MLL3-dependent histone methylation, but possibly reflect subtle genetic background effects.

### Time of day gating of inflammatory responses is not dependent on MLL3

Circadian gating mechanisms are important in immunity and host defence (Scheierman et al review). Since our analysis of the MLL3 disruption revealed changes in relation to binding sites of a number of inflammation-regulatory transcription factors with reported roles in mediating circadian control, including NR3C1, NR1D1, NR1D2, and NFIL3 (Fig 2D) we next investigated the role of MLL3 in conferring temporal control to inflammation. For this, we used *MLL3*^*f/f*^.*LysMcre* mice and *MLL3*^*f/f*^ littermate controls. Animals were exposed to lipopolysaccharide (LPS), via an intra-peritoneal route. We have previously characterised this model and shown enhanced inflammatory responses at ZT12 relative to ZT0, which is dependent on macrophages (Gibbs et al., 2012).

Analysis of circulating cytokine concentrations revealed a subtle impact with some (IL-6) showing a loss of temporal regulation in the absence of MLL3, but others (eg IL-1b) showed preserved variation (Fig 4C). Other cytokines, including TNF, GM-CSF, IL12p70, and INFg showed no time of day gating, and no significant differences between genotypes (Fig 4C). As we have recently discovered a strong circadian clock regulation of macrophage phagocytosis we also measured this by time of day in the two genotypes (Fig 4D). Here, we again saw time of day variation in phagocytosis efficiency as previously reported, but no genotype effect.

Because pulmonary inflammatory responses are strongly circadian we next moved to an aerosolised LPS model to target the lung directly. Here, we have previously characterised enhanced inflammation at ZT0 relative to ZT12, which is gated by the circadian clockwork of lung epithelial cells (Gibbs et al., 2014). Therefore we targeted the MLL3 Delta mutation specifically to lung epithelial cells using a CCSP-iCre driver (*MLL3*^*f/f*^.*CCSPiCre*). We first profiled the response to nebulised lipopolysaccharide administered at ZT0, or ZT12 in the *MLL3*^*f/f*^.*CCSP*^*icre*^ mice and *MLL3*^*f/f*^ littermate controls by assaying bronchoalveolar lavage fluid (Fig 5A). Here we saw no genotype effect on responses to bronchoalveolar lavage fluid concentration of CXCL5, IL6, or TNFa, characteristic cytokine/chemokine responsive proteins (Fig 5A). The previously reported time of day variation in CXCL5 concentration was seen in both genotypes (Gibbs et al., 2014). Furthermore, absence of MLL3 catalytic activity did not affect the circadian regulation of *Rev-erbα* by time of day, or by LPS (Fig 5 Supp A).

**Figure 5.**
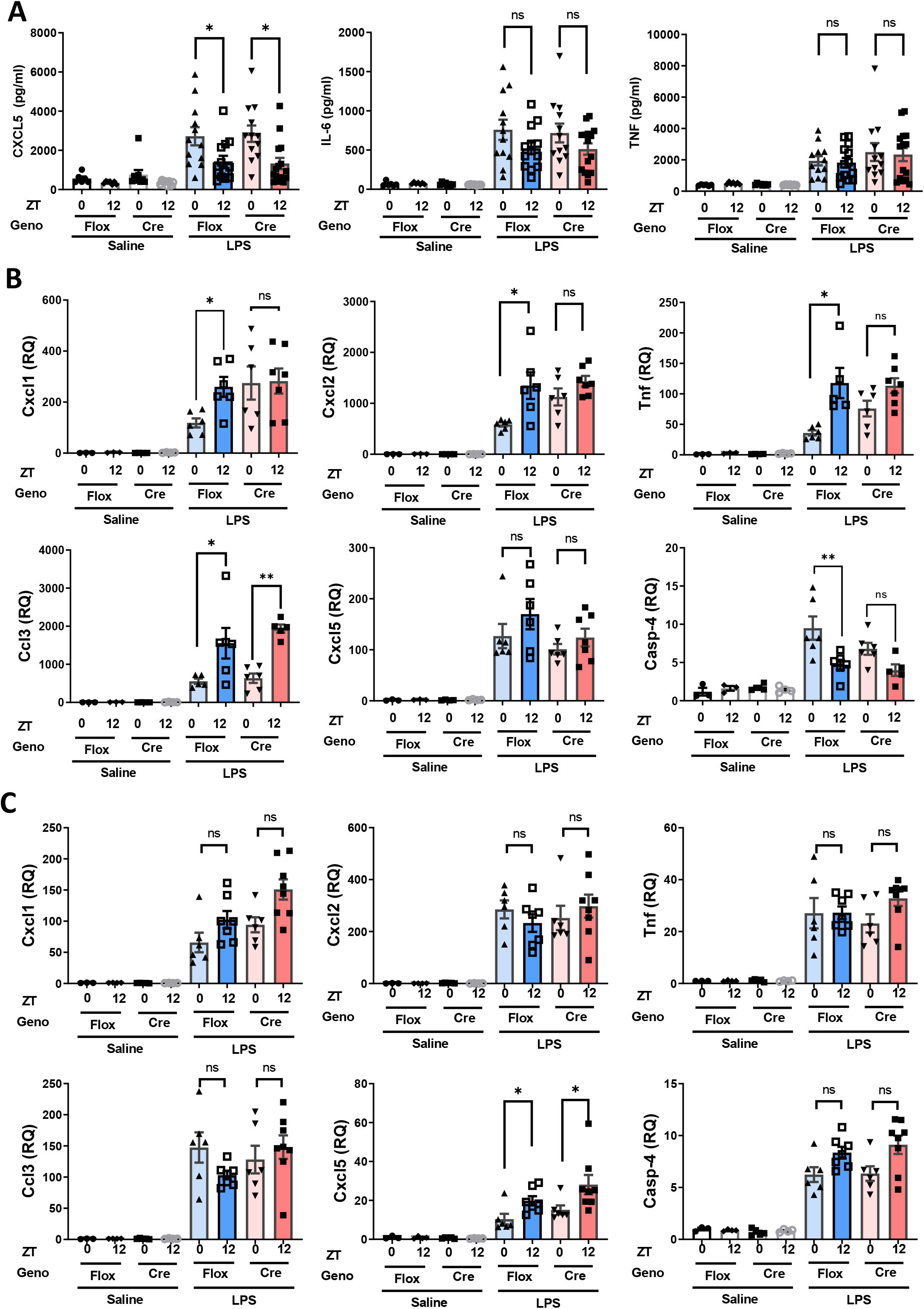
The role of MLL3 in the circadian gating of the airway inflammatory response. MLL3^Delta flox/Delta flox^ CCSPiCre^het^ mice and MLL3^Delta flox/Delta flox^ littermate controls were exposed to aerosolised LPS at ZT0 or ZT12. 5 hours after exposure bronchoalveolar lavage fluid (BALF) and lung tissue were harvested. **A**. Cytokine levels were quantified in the BALF by ELISA. mRNA levels of various genes associated with the inflammatory response were measured were quantified by qPCR. **B**. The quantified levels in male mice. **C**. The quantified levels in female mice.

In a further analysis we measured inflammatory mediator mRNA expression in lung tissue from the same animals. Here, male and female mice showed significant differences in response, and so were analysed separately (Fig 5B, C). We observed a surprising difference between the two genotypes, in males, with time of day variation in CXCL1, CXCL2, TNFa, and Caspase 4 all being lost in the absence of MLL3 (Fig 5B). The peak inflammatory chemokine expression in lung tissue is in anti-phase to that of the mature protein secreted into the airway, reflecting further complexity in the circadian regulation of integrated lung immune responses. This may in part be due to circadian regulation of the *Tlr4* receptor in the lung tissue, which was increased at ZT12 in both genotypes (Fig5 Supp B). The loss of circadian gating did not extend to all measured transcripts, with CCL3 showing a peak at ZT12 in both genotypes (Fig 5B). Intriguingly, we did not see a time of day or genotype difference in lung CXCL5 gene expression in males (Fig 5B). In females, the time of day variation in gene response was confined to CXCL5, and here we saw no impact of MLL3 loss (Fig 5C). This suggests that MLL3 is driving different biology in males and females. As MLL3 is a transcriptional co-activator, driving methylation of H3K4, and thereby promoting active promoters and enhancers it is further surprising that in the absence of MLL3 the dawn nadir of chemokine gene expression was lost. This suggests a gain in gene expression, a change perhaps best seen in CXCL1.

## Discussion

In the present study we set out to investigate the interaction between the histone methyltransferase MLL3, and the core circadian clock. The core clock TTFL controls the circadian oscillation of thousands of individual transcripts and, importantly, is able to do so in a tissue specific manner. Furthermore, the core clock gates specific cellular functions by time of day, including the inflammatory response. In order to facilitate such diverse yet fine-tuned genomic control, the core clock must coordinate with histone-modifiers, such as MLL3. Previous reports have shown that MLL3 expression is circadian, and furthermore that disruption of MLL3 catalytic activity (MLL3 Delta) in MEFs abolishes the oscillation of the core clock. Given this observation, we first investigated the effect of MLL3 Delta in mouse behavioural rhythms, but found no differences between animals lacking MLL3 methyltransferase activity, and their wild-type littermates. We further extended the studies to isolated mouse tissues and found that SCN, lung, liver, and isolated macrophages exhibited strong circadian oscillations in the absence of MLL3 catalytic activity, in stark contrast to the previously published observations in MEFs (Valekunja et al., 2013). However, we did observe tissue-specific changes in period in liver and peritoneal macrophages in mice with global MLL3 Delta mutation. To investigate the apparent disconnect between our observations in mature tissues, and those previously published in MEFs, we performed H3K4me1 and H3K4me3 ChIP-seq in liver tissue from MLL3 Delta mice and littermate wild-type controls. We found surprisingly subtle changes in genomic histone methylation, explaining the lack of circadian phenotype, and implying compensation from other methyltransferases. Finally, we examined the potential role of MLL3 as an effector of circadian time-of-day gating of the inflammatory response, where we identified subtle and potentially sexually dimorphic roles for MLL3. Our findings require a major reassessment of the role of MLL3 in circadian oscillator function.

Given the strong effect of MLL3 Delta on development, we were surprised to find only subtle effects on the circadian clock. It has been previously reported that the MLL3 Delta mutation is embryo lethal for homozygotes on a pure C57Bl6 genetic background (Lee, 2006). We too found this to be the case, and therefore bred global MLL3 Delta mice expressing Per2::luc on a mixed C57Bl6/129sv genetic background. Even on this mixed genetic background we found a significant deviation from the expected Mendelian ratio, with fewer viable homozygous MLL3 Delta mice born (Fig1 S1). However, in the surviving MLL3 Delta homozygotes, no effect on circadian behavioural parameters was observed, and in corroboration, oscillations in isolated SCN were equivalent between Delta mice and wild-type littermate controls. Intriguingly, we observed a significant shortening of circadian period in other isolated tissues, specifically liver and peritoneal macrophages. This indicated that there may be some subtle interactions between MLL3-dependent methylation and core clock function in a tissue-specific manner, however it is difficult to reconcile these observations with the stark and complete loss of oscillation reported in MEFs (Valekunja et al., 2013).

In order to investigate the difference between our observations and those in MEFs, we performed ChIP-seq. Valekunja et al found circadian oscillations in H3K4me3 deposition and MLL3-genome association in mouse liver, peaking at CT18, and complete loss of global H3K4me3 in MEFs from MLL3 Delta mice. We therefore harvested liver tissue from MLL3 Delta mice and wild-type littermate controls at ZT18 and performed H3K4me3 ChIP-seq. Again, in stark contrast to the findings of Valekunja et al, we found no difference in the overall global levels of H3K4me3 deposition between the two genotypes, and only subtle repositioning of the mark in the Delta condition. Furthermore, we found no differences in the H3K4me3 deposition across the promoters of the core clock genes, explaining why a fully functional circadian clock was observed in MLL3 Delta liver tissue. Our observations are in line with other reports that find little impact of MLL3 catalytic inactivation on gene expression (Dorighi et al., 2017; Rickels et al., 2020, 2017). This may be due to functional redundancy with MLL4 (Lee et al., 2008, 2009)

Detailed reports have found that MLL3 has considerably stronger activity depositing H3K4me1, and that H3K4me3 is mostly deposited by other methyltransferases, such as MLL1 (Katada and Sassone-Corsi, 2010). We therefore also performed H3K4me1 ChIP-seq in liver, as direct effects of MLL3 enzymatic disruption may be more easily observed. H3K4me1 denotes enhancers, which modulate the expression of genes from a range of thousands to millions of basepairs away. For this reason it is difficult to assign changes in enhancer H3K4me1 deposition to changes in gene expression, and examining the H3K4me1 levels proximal to genes of interest is not necessarily informative. Regardless, little difference in H3K4me1 levels was found in the loci of core clock genes. Furthermore, the global levels of H3K4me1 deposition were comparable between the two genotypes and, as with H3K4me3, only subtle repositioning of H3K4me1 peaks was observed. Interestingly, globally, there was a redistribution of H3K4me1 peaks from intronic regions to intergenic and promoter regions in the MLL3 Delta liver tissue compared to the wild-type littermate controls (Figure 3D). This may be due to a compensation of other histone methyltransferases with a selectivity for different genomic regions/substrates to that of MLL3. Indeed, GIGGLE score analysis identified an overlap of MLL3 Delta specific H3K4me1 peaks with previously published H2A.Z ChIP-seq peak locations, which may indicate that in the absence of MLL3 catalytic activity the COMPASS complex is being drawn to more active chromatin.

Having found that the oscillation of the core clock was intact in MLL3 Delta mice we next examined whether MLL3 methyltransferase activity may confer time-of-day control to known circadian-regulated processes. We chose to examine the inflammatory response, which we have previously extensively characterised, is strongly circadian (Baxter and Ray, 2019; Gibbs et al., 2014, 2012; Pariollaud et al., 2018). We examined two models of TLR4-driven inflammation, the first a systemic response to IP injection of LPS, which we have previously shown is gated by the macrophage cell clock, and secondly aerosolised LPS, which shows a circadian response that is gated by epithelial cell clocks. Interestingly, the enhancement of inflammation in these two models is in anti-phase: enhanced inflammatory response in the systemic model is observed at ZT12, whereas enhanced inflammatory response in the aerosolised model is observed at ZT0, indicating different circadian mechanisms.

In each case we targeted the MLL3 Delta mutation to the specific cell type which confers circadian control to the inflammatory response using Cre-driven recombination of MLL3 exons 57-58.

One of the key cytokines driving the inflammatory response in the systemic model (IP LPS) is IL-6, and a higher IL-6 response was found at ZT12 relative to ZT0 in wild-types. Time of day gating of IL-6 was lost in the MLL3^flox^.LysMcre mice, however only subtle differenes were seen in the circulating concentrations of other cytokines. By contrast, the circadian gating of the airway response to aerosolised LPS is driven by CXCL5 secretion (Gibbs et al., 2014). This time of day gating was maintained in MLL3^flox^.CCSPiCre mice. Taken together this indicates a specificity of MLL3 action in conferring circadian control. In the aerosolised LPS model, we went on to look at the mRNA induction in the lung tissue. Surprisingly we found that the circadian peak of many inflammatory genes was in anti-phase to the peak concentration of cytokines secreted into the airway lumen. In the lung tissue we found a sexually dimorphic inflammatory response, in line with previous publications (Marriott et al., 2006; Merkel et al., 2001). Furthermore, we found a sex-specific role for MLL3 in the gating of the inflammatory response, for example in the case of *Cxcl1* and *Tnf*.

Taken together, we find that MLL3 is not required for circadian cycling of core clock genes, in strong contrast to previous reports. We do, however, find subtle, sex-specific effects of MLL3 on H3K4me1 distribution across the genome, and on circadian gating of the inflammatory response in peripheral tissues. Further investigations into the interaction of histone modifying complexes and the circadian clock is warranted, and may lead to better understanding of the inflammatory response in relation to time-of-day specificity, and sexual dimorphism.

## Acknowledgements

The work is supported by an Wellcome Investigator Award. DWR and ASIL are Wellcome Investigators, Wellcome Trust (107849/Z/15/Z, 107851/Z/15/2. We would like to thank Professor Jae W. Lee for kindly providing the MLL3 Delta mouse strain. We would like to thank Professor Joseph S. Takahashi for kindly providing the Per2::LUC mouse strain.

## Author contributions

MB, ASIL, and DWR designed the study. MB, TP, PC, LH, MV, GK, LG, NB, DK, AH, and RM performed data collection and/or bioinformatic analysis. MB and DWR wrote the manuscript. ASIL, PC, LH, RM, and MV supported manuscript preparation.

## Figure Legends

**Figure 1. Supplement.**
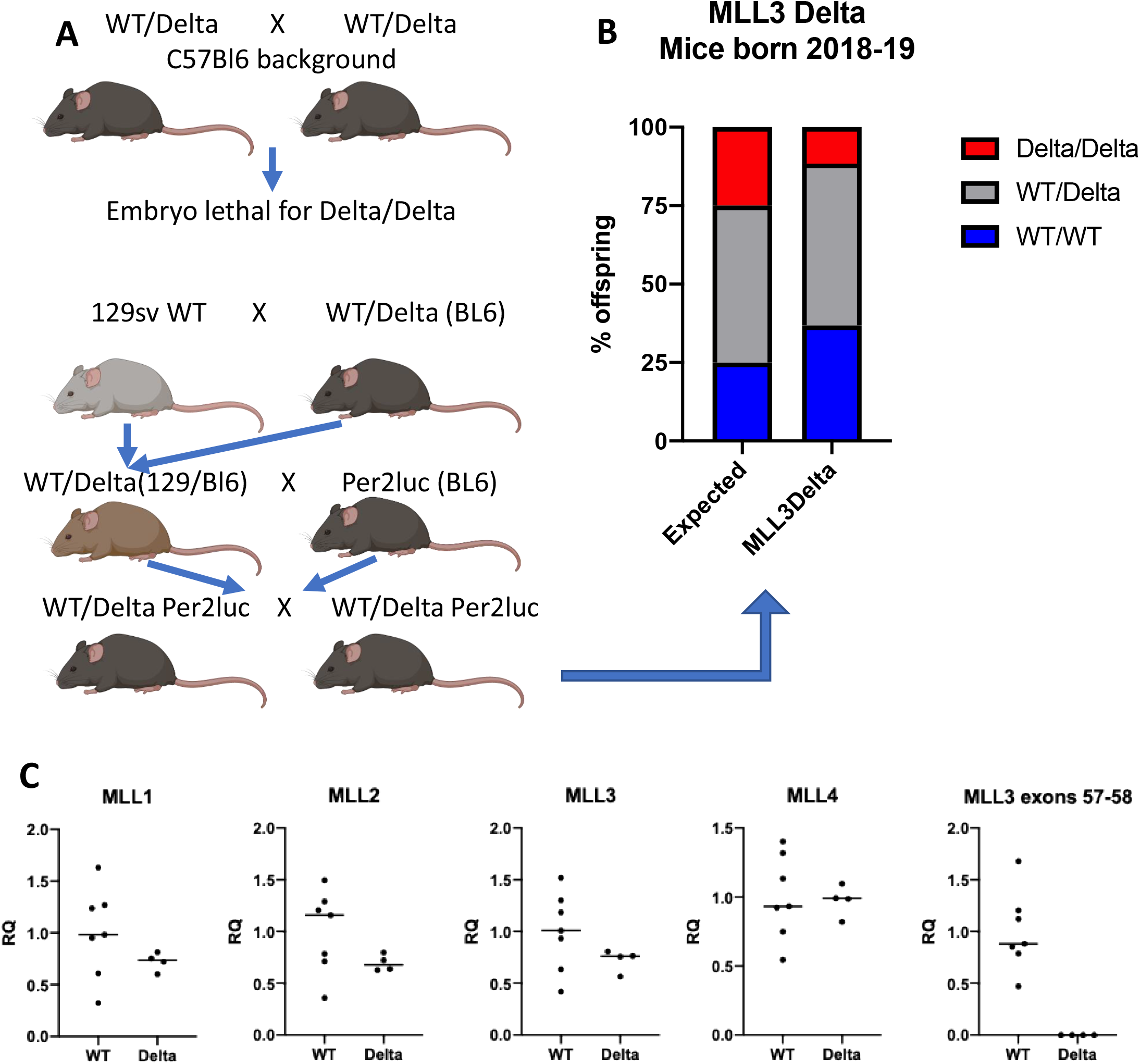
**A**. Breeding scheme to show the development of MLL3 Delta and wild-type control mice on a mixed C57Bl6/129s background, expressing Per2::luc. **B**. quantification of viable mice born from MLL3 Delta heterozygous crosses on C57Bl6/129s background. The number of mice which were homozygous, heterozygous and wild-type for the MLL3 Delta mutation is shown, alongside the expected Mendelian ratio. **C**. mRNA expression analysis of MLL3 alleles and other MLL family members which may form part of the COMPASS complex, from MLL3 Delta animals and wild-type controls.

**Figure 2. Supplement.**
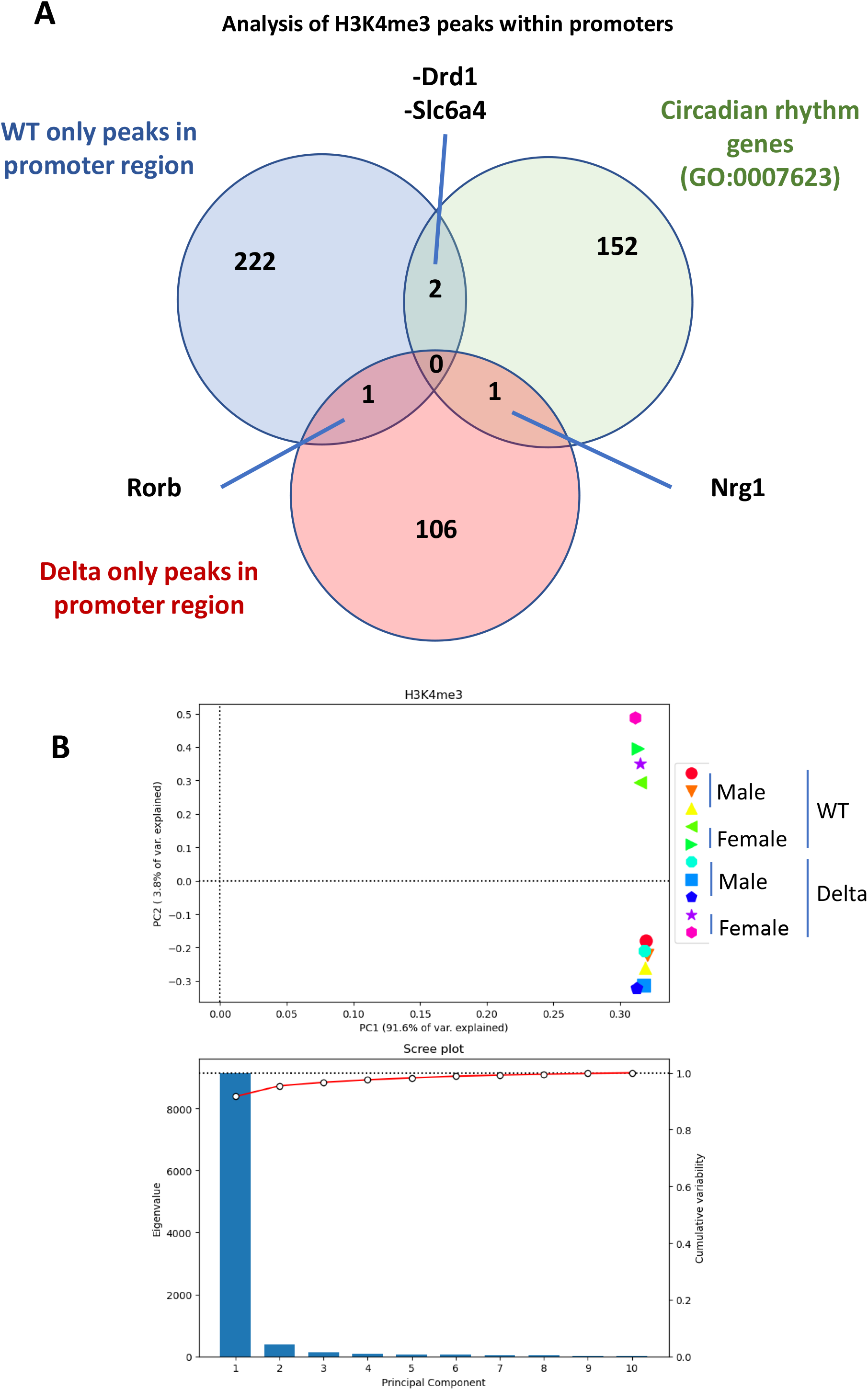
Analysis of positioning of H3K4me3 peaks which were unique to either the WT or Delta condition. Only peaks within gene promoters were analysed. The Venn diagram shows the number of genes with a unique H3K4me3 peak in their promoter in either the WT or Delta condition. These genes are further compared with genes that fall into the “Circadian rhythm genes” GO category. PCA analysis of all H3K4me3 ChIP-seq samples using Deeptools.

**Figure 3 Supplement.**
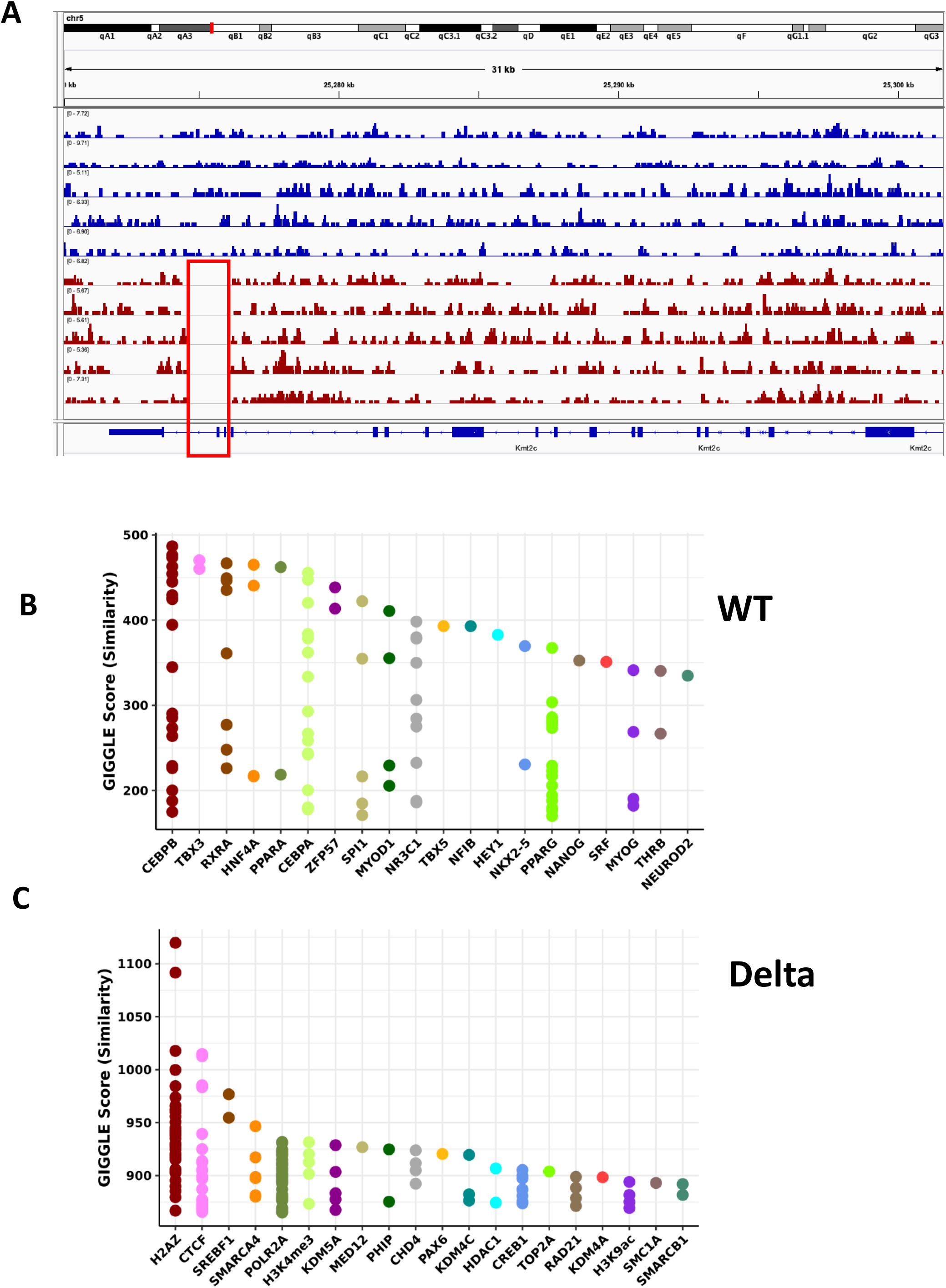
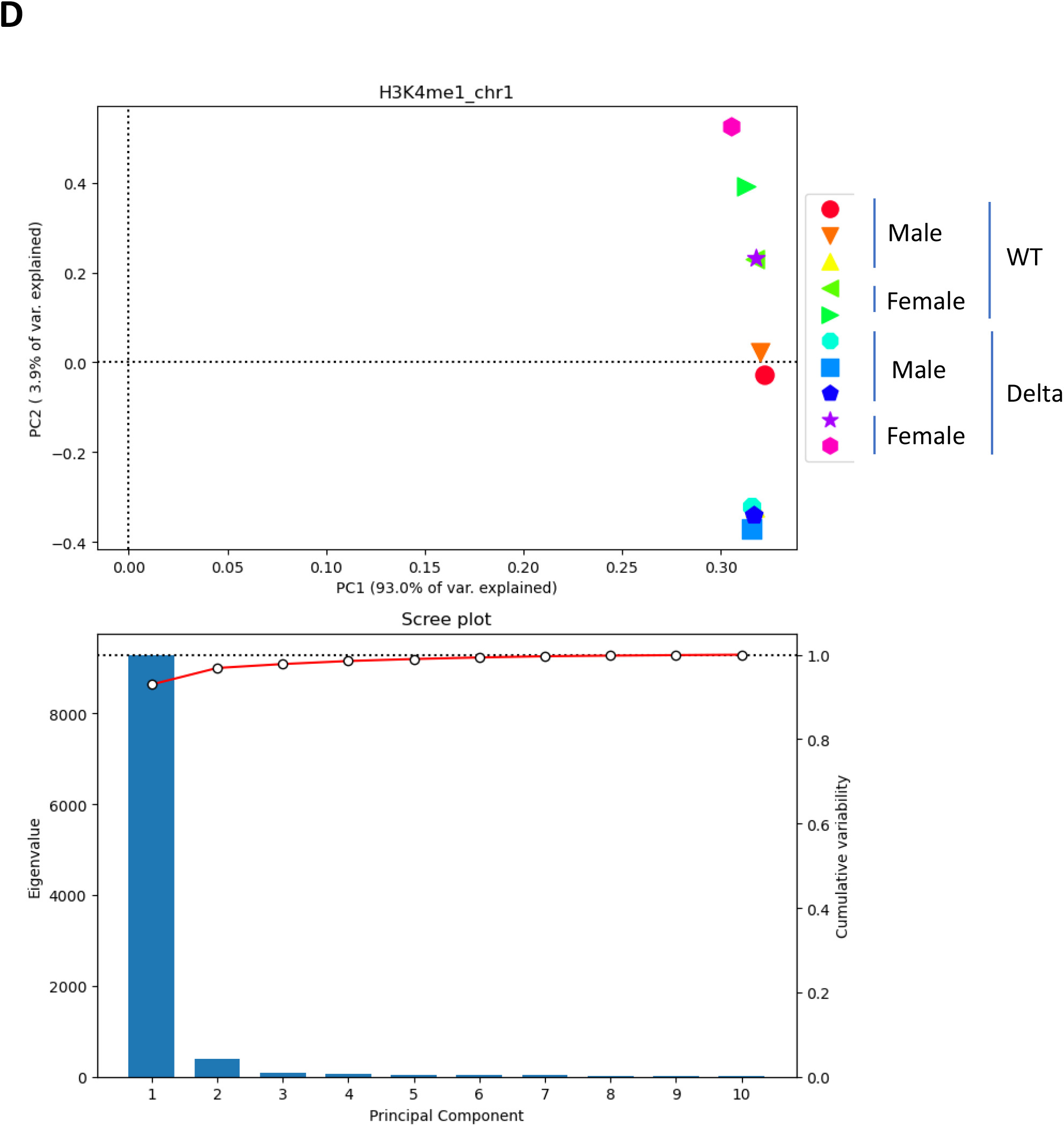
**A**. Gene tracks for each individual mouse across the MLL3 genomic locus. The deleted region of the MLL3 Delta allele, across exons 57-58, is highlighted in the red box. Tracks from wild-type mice are in blue, tracks from Delta mice are in red. **B**. H3K4me1 peaks which were found to be unique to the WT condition were submitted for Giggle score analysis to assess similarity with other published ChIP-seq factor binding datasets. **C**. Peaks which were found to be unique to the Delta condition were submitted for Giggle score analysis to assess similarity with other published ChIP-seq factor binding datasets. **D**. PCA analysis of all H3K4me3 ChIP-seq samples using Deeptools.

**Figure 4 Supplement.**
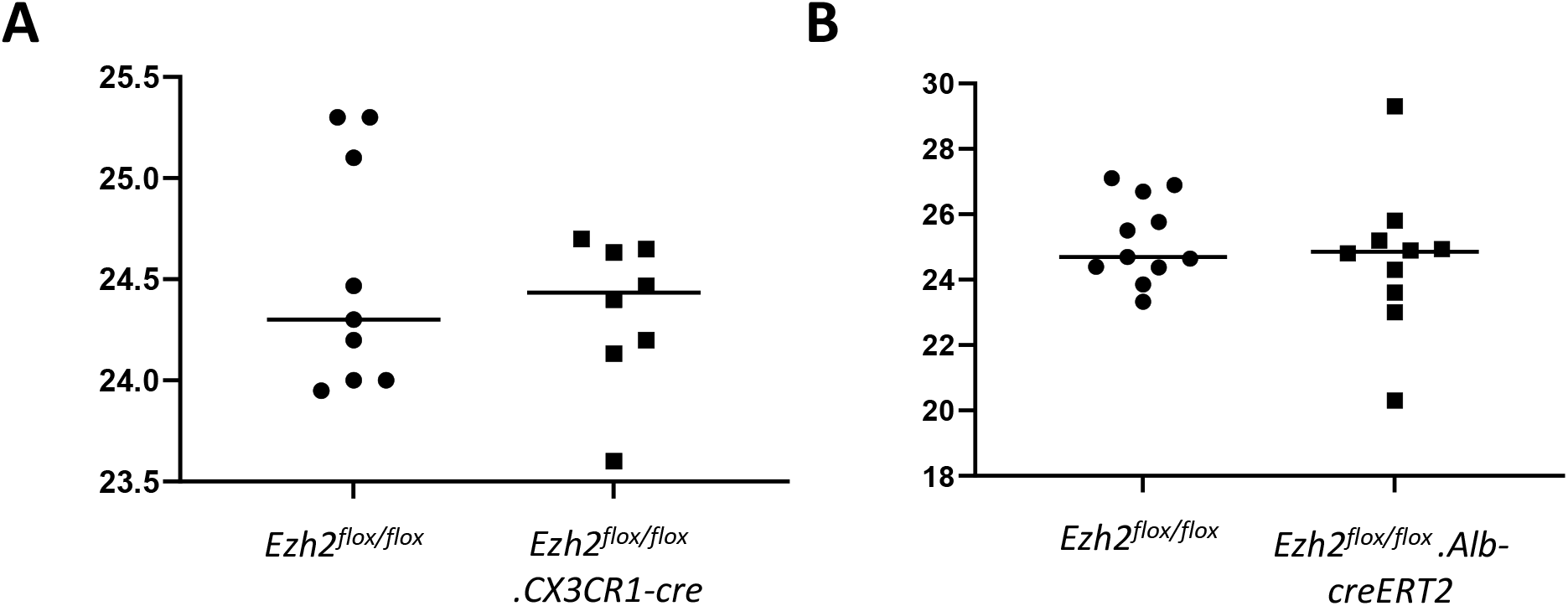
**A**. PECs were isolated from Ezh2 floxed Cx3cr1-Cre mice and Ezh2 floxed littermate controls, and placed into a lumicycle. Circadian oscillations in Per2::luc were measured for 3 days and period was calculated. Each data point represents the average of 3 technical replicates from an individual animal. **B**. Liver slices were taken from Ezh2 floxed AlbCreERT2 mice and Ezh2 floxed littermate controls, treated with tamoxifen for 5 days. Liver slices were taken 10 days after the final tamoxifen treatment and placed in a lumicycle. Circadian oscillations in Per2::luc were measured for 3 days and period was calculated. Each data point represents the average of 3 technical replicates from an individual animal.

**Figure 5 Supplement.**
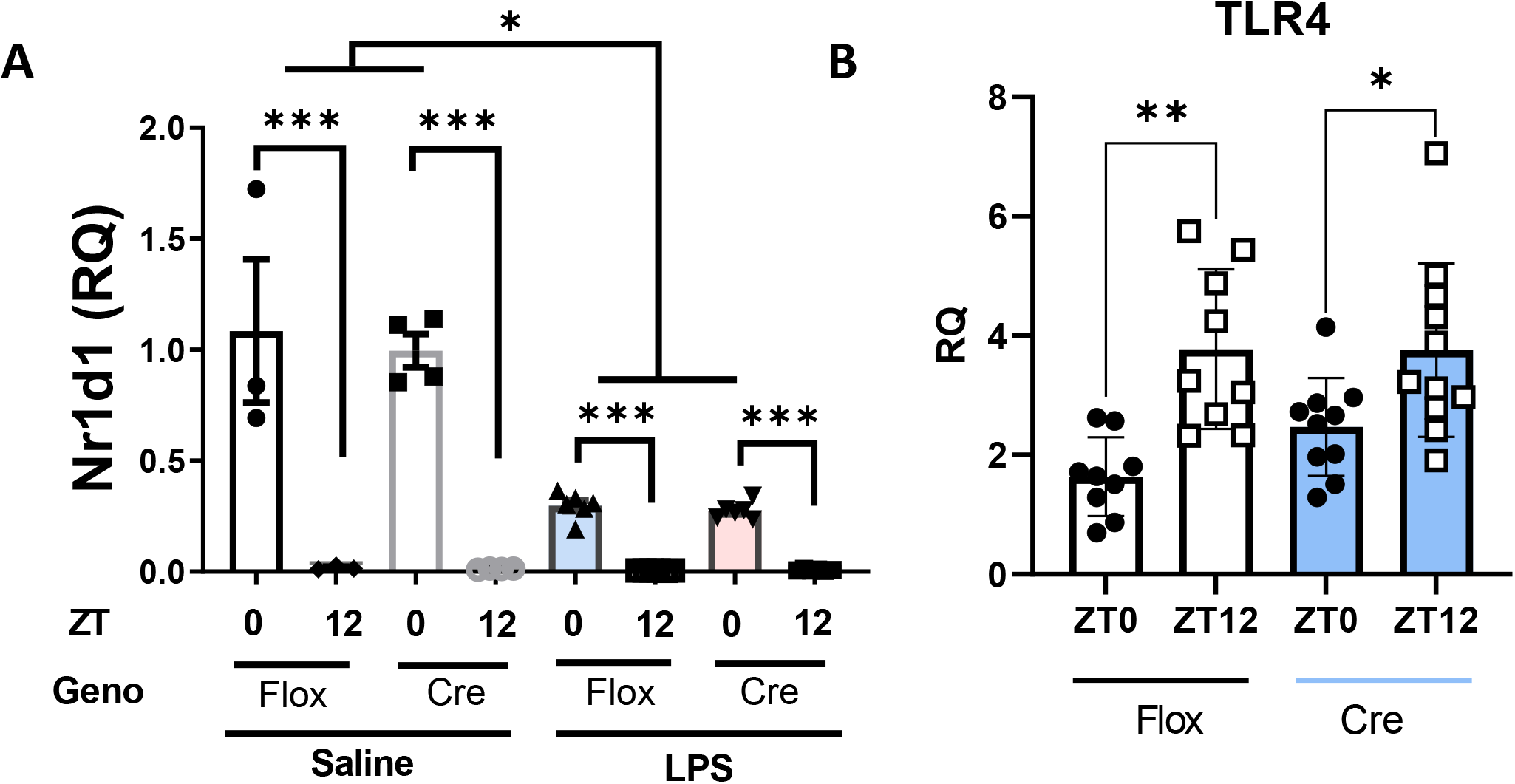
**A**. qPCR analysis of *Reverba (NR1D1)* expression in lung tissue from mice exposed to aerosolised LPS or saline control at ZT0 or ZT12. Lung tissue was harvested 5 hours after exposure. **B**. qPCR analysis of *Tlr4* expression in lung tissue from mice exposed to aerosolised LPS or saline control at ZT0 or ZT12. Lung tissue was harvested 5 hours after exposure. As there was no effect of LPS exposure on *Tlr4* expression, saline and LPS exposed data were pooled by timepoint and genotype.

## Materials and Methods

### Animals

All mice were routinely housed in 12:12 light/dark (L:D) cycles, unless otherwise stated, in light-controlled chambers, with *ad libitum* access to food and water. Light exposure within the chambers was continuously monitored throughout experiments. All experiments were carried out in accordance with the Animals (Scientific Procedures) Act 1986 (UK). MLL3 Delta mice and MLL3^fl/fl^ mice were kindly provided by Professor Jae W. Lee. MLL3 Delta mice have a global in-frame deletion of *Mll3* exons 57-58. Mice were bred as heterozygous pairs, producing mice homozygous for the Delta allele (MLL3 Delta mice), and animals homozygous for wild-type MLL3 (wild-type littermate controls). MLL3^fl/fl^ mice have exons 57-58 floxed, allowing recombination through cell-type specific Cre expression. For experiments using conditional targeted mice, experimental animals were heterozygous for Cre expression, and controls were littermates (floxed/floxed) carrying no copies of the Cre. mPer2::luc mice were kindly provided by Professor Joe Takahashi (Yoo et al., 2004). Locomotor activity was recorded in home cages using infrared beam-break sensors, as previously described (Cunningham et al., 2016). Genotyping was performed on all animals, using previously published protocols (Clausen et al., 1999; Lee et al., 2006; Li et al., 2012).

## Bioluminescence Recording

For *ex vivo* analyses of circadian rhythms, mice were bred onto a mPER2:luc genetic background, and tissues were harvested between 8-12 weeks of age. mPER2-dependent bioluminescence was recorded from liver, lung, SCN slices, or cultured peritoneal macrophages (PECs), all maintained at 37 °C using a Lumicycle (Actimetrics) as described previously (Meng et al., 2010). Circadian period was measured from the 2^nd^ peak post-culture for two complete cycles. 2-3 independent tissue samples were averaged for each animal. WDR5-specific compound OICR9429 was purchased from Tocris, and made up in DMSO. PECs were treated with OICR9429 at specified concentrations, with 0.1% DMSO final concentration in DMEM/F12 media.

### Systemic LPS

Mice were challenged with intraperitoneal (IP) lipopolysaccharide (LPS; 1mg/kg; isolated from E.Coli 0127:B8, L4516, Sigma), at either ZT0 or ZT12, as previously described (Gibbs et al., 2012). Blood plasma was isolated 5 hours after challenge. Cytokine analysis in plasma was performed by ELISA (R&D Systems), or Magpix assay (Luminex).

### Aerosolised LPS

Mice were exposed to aerosolised LPS (2 mg/mL; isolated from E.Coli 0127:B8, L4516, Sigma) or saline (0.9%) for 20 minutes, as previously described (Gibbs et al., 2014). Mice were euthanised 5 hours after exposure. Bronchoalveolar lavage fluid was collected for cytokine secretion analysis by ELISA (R&D Systems). Lung tissue was collected for gene expression analysis by qPCR (SYBR Green, see table for primer sequences).

### ChIP-seq

Liver tissue was collected from MLL3 Delta mice and littermate wild-type controls at ZT18. ChIP reactions were performed by Active Motif (Carlsbad, USA), using antibodies AB_2615075 and AB_2615077 for H3K4me1 and H3K4me3 respectively. The ChIP reactions also contained drosophila chromatin spike-in for normalisation of sequencing data. ChIP-seq libraries were generated from the ChIP-and Input-DNA using a custom Illumina library type on an automated system (Apollo 342, Wafergen Biosystems/Takara). ChIP-Seq libraries were sequenced on Illumina NextSeq 500 as 75-nt single end reads.

### Data processing

Data was processed as previously published (Hunter et al., 2020). QC on FastQ files was performed using FastQC v0.11.8 (Wingett and Andrews, 2018). Trimmomatic v0.39 (Bolger et al., 2014) was used to trim adapters and remove poor quality reads: *java -jar trimmomatic-0*.*39*.*jar SE -phred33 <FILENAME*.*fastq> ILLUMINACLIP:TruSeq3-SE*.*fa:2:30:10 LEADING:3 TRAILING:3 SLIDINGWINDOW:4:15 MINLEN:36*.

Trimmed reads were aligned to the mm10 reference genome using Bowtie2 v2.4.1 (Langmead and Salzberg, 2012): *bowtie2 -p 6 -x mm10 -U <FILENAME*.*fastq> -S <OUTPUT_FILENAME*.*sam*) SAMtools v1.10 (Li et al., 2009) was used to produce BAM files from SAM files (*view, sort, index*, all with default settings. Picard v2.22.9 was used to remove duplicate reads: *java -jar picard*.*jar MarkDuplicates I=<FILENAME*.*bam> O=<OUTPUT_FILENMAE*.*bam> M=<METRICS_FILENAME*.*txt> REMOVE_DUPLICATES=true ASSUME_SORTED=true VALIDATION_STRINGENCY=LENIENT USE_JDK_DEFLATER=true*

### Peak Calling

ChIP-seq peaks were called in processed BAM files against processed control (input) files using MACS2 v2.2.7.1 (Zhang et al., 2008). Peaks were called using the following parameters: *Macs2 callpeak -t <SAMPLENAMES*.*bam> c <INPUT*.*bam> --name <OUTPUT_FILENAME> -f BAM -g mm --keep-dup=1 -q 0*.*01 --bdg --SPMR --verbose 0 > <OUTPUTFILENAME* The narrowpeak setting was used to call peaks in the H3K4me3 data, whilst the broadpeak setting was used for H3K4me1 data (*--broad*). Peaks were counted using the respective commands: *wc -l <FILENAME*.*narrowpeak> narrowPeak wc -l <FILENAME*.*broadpeak> broadPeak*BEDtools v2.29.2 (Quinlan, 2014) was used to find peaks which were common between datasets: *bedtools intersect -u -a <FILENAME*.*broadPeak> -b FILENAME2*.*broadPeak > <OUTPUT_FILENAME*.*bed*>

### ChIP-seq Data Visualisation

deepTools v3.5.1 (Ramírez et al., 2014) was used to make heatmaps of signal intensity, using the commands *computeMatrix* and *plotHeatmap*. Integrative Genomics Viewer v2.9.4 (Thorvaldsdóttir et al., 2013) was used to produce visualisations of ChIP-seq data tracks.

### Statistics

Statistical tests and sample numbers are specified in figure legends where appropriate. Unless otherwise stated, statistical significance was tested using Mann-Whitney U test. Statistical tests were conducted in GraphPad Prism. Throughout, * denotes p<0.05, ** denotes p<0.01, *** denotes p<0.001 and Plots were produced in GraphPad Prism, except for ChIP-seq data, which is described above.

### Data Availability statement

ChIP-seq data has been uploaded to ArrayExpress. Accession number E-MTAB-1120.

